# Enforced dimerization of CD45 by the adenovirus E3/49K protein inhibits T cell receptor signaling

**DOI:** 10.1101/2023.02.23.529489

**Authors:** Mark Windheim, Thomas F. Reubold, Matthias Gaestel, Hans-Gerhard Burgert

## Abstract

Human adenoviruses (HAdVs) are widespread pathogens that generally cause mild infections in immunocompetent individuals, but severe or even fatal diseases in immunocompromised patients. In order to counteract the host immune defenses HAdVs encode various immunomodulatory proteins in the early transcription unit 3 (E3). The E3/49K protein is a highly glycosylated type I transmembrane protein uniquely expressed by species D adenoviruses. Its N-terminal ectodomain sec49K is released by metalloprotease-mediated shedding at the cell surface and binds to the receptor-like protein tyrosine phosphatase CD45, a critical regulator of leukocyte activation and functions. It remained elusive which domains of CD45 and E3/49K are involved in the interaction and whether such an interaction can also occur on the cell surface with membrane-anchored full-length E3/49K. Here, we show that the two extracellular domains R1 and R2 of E3/49K bind to the same site in the domain d3 of CD45. This interaction enforces the dimerization of CD45 causing the inhibition of T cell receptor signaling. Intriguingly, the membrane-anchored E3/49K appears to be designed like a “molecular fishing rod” using an extended disordered region of E3/49K as a “fishing line” to bridge the distance between the plasma membrane of infected cells and the CD45 binding site on T cells to effectively position the domains R1 and R2 as baits for CD45 binding. This design strongly suggests that both the secreted sec49K and the membrane-anchored full-length E3/49K have immunomodulatory functions. The forced dimerization of CD45 may be applied as a therapeutic strategy in chronic inflammatory disorders and cancer.

## Introduction

Human adenoviruses (HAdVs) typically induce self-limiting infections associated with acute disease in immunocompetent individuals, but in immunocompromised patients HAdV infections tend to be severe or even fatal (Lion, 2019; Wold & Ison, 2013). There are more than 100 HAdV types that are classified into species A-G (Berk, 2013; Robinson et al., 2013) and the great majority (>70 types) belong to species D (Ismail et al., 2018; Ismail et al., 2019; Robinson et al., 2011). Interestingly, only a few of these HAdV-D types, namely HAdV-D8, 37, 53, 54, 56 and 64 (formerly Ad19a) cause epidemic keratoconjunctivitis (EKC), a highly contagious and severe eye disease involving both the conjunctiva and the cornea (Ismail et al., 2019; Kuo, 2019; Rajaiya et al., 2021). HAdVs have also been exploited as vectors for gene therapy, cancer therapy and vaccination, e.g. against SARS-CoV-2/COVID-19 (Gao et al., 2020; Jacob-Dolan & Barouch, 2022; Watanabe et al., 2021). In these vectors the early transcription unit 3 (E3) is typically deleted to create space for an insert, because the E3 region is not required for HAdV replication *in vitro* (Berk, 2013; Wold & Ison, 2013). Nevertheless, the E3 region is preserved in all HAdVs, indicating an important role *in vivo.* Many functional investigations *in vitro* showed that E3 proteins counteract host immune responses (Burgert & Blusch, 2000; Burgert et al., 2002; Oliveira & Bouvier, 2019). Species D HAdVs have the largest E3 region encoding eight open reading frames (ORFs). One of these genes, designated E3/49K (or CR1-β), is only present in species D HAdVs (Blusch et al., 2002; Windheim & Burgert, 2002; Windheim et al., 2004).

The E3/49K ORF was initially identified in the E3 region of the epidemic keratoconjunctivitis (EKC)-causing HAdV-D64 (Deryckere & Burgert, 1996). E3/49K is a highly glycosylated type I transmembrane protein with a short cytoplasmic tail and an extracellular domain (ECD) comprising three internal repeats designated conserved regions 1-3 (R1-3) that are predicted to form immunoglobulin-like domains (Blusch et al., 2002; Windheim & Burgert, 2002). E3/49K is synthesized in the early phase of infection, but continues to be produced in the late phase (Windheim & Burgert, 2002). Interestingly, E3/49K is cleaved by a matrix metalloprotease (MMP) N-terminal to the transmembrane domain, generating a small membrane-integrated 12-14 kDa C-terminal fragment and a large secreted ectodomain, named sec49K (Windheim et al., 2016; Windheim et al., 2013). Sec49K is the only secreted HAdV protein known to date. Functionally, we demonstrated that sec49K binds to the cell surface receptor-like protein tyrosine phosphatase CD45 on all leukocytes, impairing e.g. the activation of CD4^+^ T cells and NK cells, thereby inhibiting cytokine production (IFN-γ/TNFα) and cytotoxicity, respectively (Windheim et al., 2013). More recently, other species D E3/49K proteins have been shown to bind to CD45 with high affinity (Martinez-Martin et al., 2016).

Interestingly, there is a homology between E3 proteins and members of the human cytomegalovirus (HCMV) RL11 gene family (Davison et al., 2003). One of these proteins, the HCMV UL11 protein, was also shown to bind to CD45 to reduce T cell signaling and proliferation and to increase IL-10 secretion as a recombinant Fc fusion protein (Gabaev et al., 2011; Osanyinlusi et al., 2022; Zischke et al., 2017). Conspicuously, CD45 is also targeted in infected cells by the m42 protein of murine cytomegalovirus (MCMV) that down-regulates cell surface expression of CD45 in RAW264.7 macrophages by triggering its lysosomal degradation (Thiel et al., 2016) and by the T-cell-tropic roseoloviruses which downregulate CD45 transcripts (Whyte et al., 2021).

CD45 is a receptor-like protein tyrosine phosphatase, which is highly expressed on all leukocytes and constitutes up to 10% of the total surface protein (Thomas, 1989). Due to differential splicing of exons 4, 5 and 6, several isoforms of CD45 are expressed and differ in the ECD with regard to the presence of regions A, B and C (Hermiston et al., 2003; Rheinlander et al., 2018; Trowbridge & Thomas, 1994). By contrast, the membrane-proximal extracellular domains d1-d4 and the intracellular tandem protein tyrosine phosphatase domains, D1 and D2, are present in all CD45 isoforms (Chang et al., 2016). Only the phosphatase domain D1 is catalytically active (Nam et al., 2005; Streuli et al., 1990). The extracellular domains of CD45 are highly glycosylated with N-glycans primarily found in the membrane-proximal domains d1-d4 and O-glycans in the membrane-distal A, B and C regions (Earl & Baum, 2008). Lectins like galectins, CD22, mannose receptor or the placental protein 14 (PP14) can bind to CD45 (Ish-Shalom et al., 2006; Martinez-Pomares et al., 1999; Symons et al., 2000; van der Merwe et al., 1996). But to date, no physiological extracellular ligand for the large proteinaceous part of the CD45 ECD has been identified.

The function of CD45 has been investigated mainly in T cells, where it is essential for activation and proliferation (Koretzky et al., 1990; Pingel & Thomas, 1989). CD45 is prominently positioned at the top of the signaling cascades originating from the T cell receptor (TCR) (Courtney et al., 2018; Gaud et al., 2018). Signaling downstream of the TCR is initiated by the dephosphorylation of the inhibitory pY505 near the C-terminus of the Src-family protein tyrosine kinase Lck (Sieh et al., 1993), which is followed by the activating phosphorylation of Y394 for maximal Lck activity (Philipsen et al., 2017). However, also the activating pY394 is a substrate of CD45 phosphatase activity (D’Oro & Ashwell, 1999), which lead to the concept that CD45 functions as a rheostat of Lck and T cell signaling (McNeill et al., 2007).

In this study, we identified the domains in CD45 and E3/49K involved in the interaction and determined the stoichiometry of the complex. We identified two CD45 binding sites in E3/49K and one E3/49K binding site in CD45 triggering the E3/49K-mediated dimerization of CD45. We propose that E3/49K enforces the dimerization of CD45 to position the intracellular phosphatase domains in such a way that access of substrates to the active site is sterically hindered. As a result, the dimerization-induced inhibition of CD45 phosphatase activity results in impaired T cell receptor signaling.

## Results

### Sec49K inhibits Lck dephosphorylation and TCR signaling

For functional studies, we previously purified the secreted form of E3/49K, sec49K, from the supernatant of cells stably expressing the full-length 49K protein (Windheim et al., 2013). However, for detailed interaction and structural studies the yield of sec49K obtained in this way was insufficient. When expressed in bacteria, the N-terminal part of 49K was largely insoluble and precipitated in inclusion bodies (data not shown). To overcome this problem, we expressed recombinant sec49K using baculovirus in insect cells, which provide N-glycosylation, albeit in a simpler way than mammalian cells (van Oers et al., 2015). To facilitate expression and detection, we added a melittin signal sequence and a C-terminal 6xHis-tag to a construct encoding residues 20-353 of 49K and inserted this into a baculovirus genome for expression in Sf (Spodoptera frugiperda) 9 insect cells. This His-tagged 49K protein (sec49K-His) could then be purified from the supernatant of infected Sf9 cells by Ni-NTA affinity chromatography. Purified sec49K-His bound with high affinity to human Jurkat T cells (Fig. 1A), as previously shown for native sec49K purified from cells expressing the full-length 49K protein (Windheim et al., 2013).

**Figure 1.**
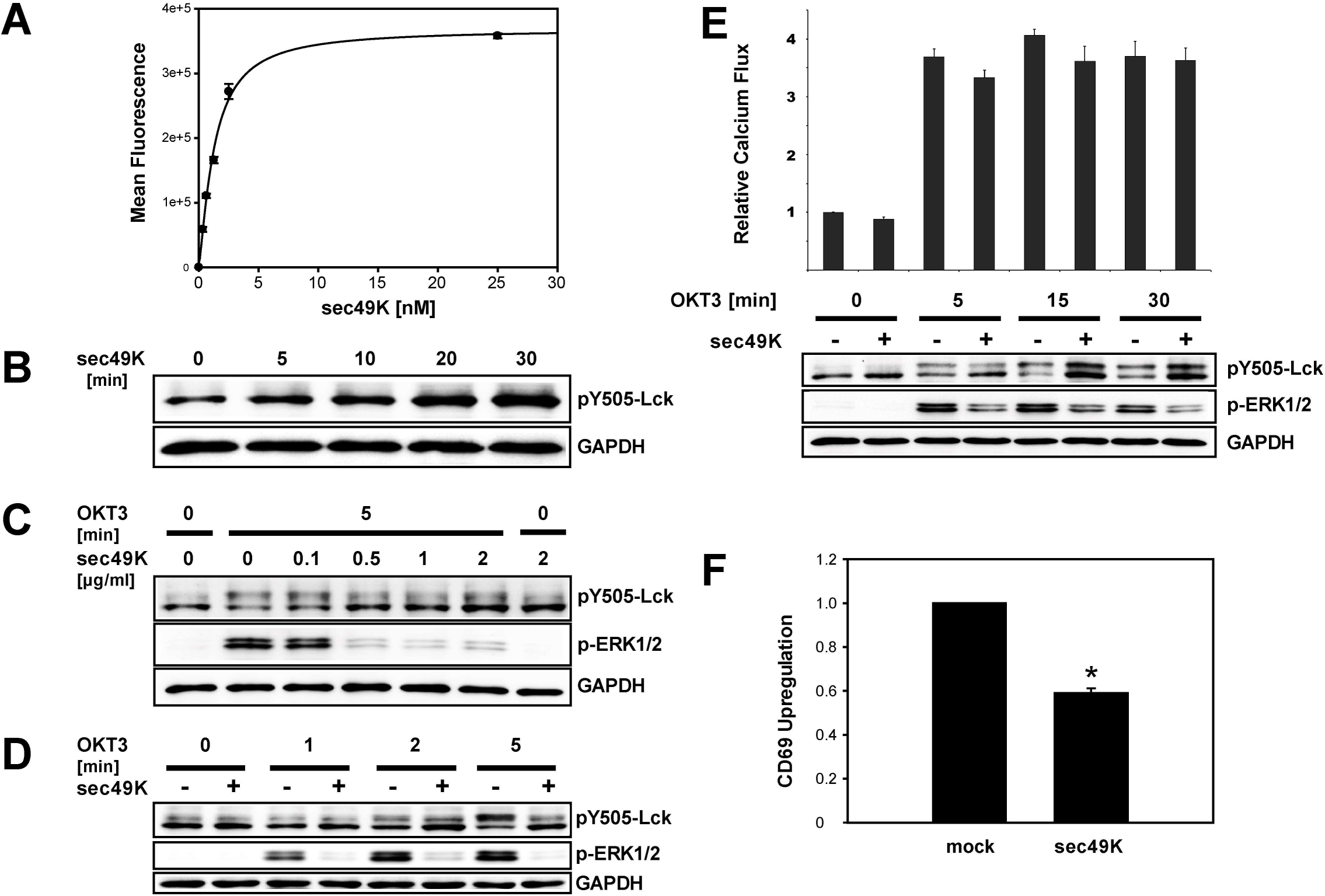
Sec49K inhibits Lck dephosphorylation and TCR signaling. (**A**) Binding of sec49K-His to Jurkat T cells was analyzed by flow cytometry after incubation of cells with different concentrations of sec49K-His as indicated and subsequent detection of sec49K-His with an anti-49K ECD rabbit antiserum (R48). The results are representative of at least three independent experiments. The error bars indicate the standard deviation between duplicate measurements. (**B**) Serum-starved Jurkat T cells were incubated with 1 μg/ml sec49K-His and lysed at different time points as indicated. Cell extracts were analyzed by immunoblotting with antibodies recognizing the proteins indicated. The results are representative of three independent experiments. (**C**) Serum-starved Jurkat T cells were incubated for 30 min with different concentrations of sec49K-His as indicated. Subsequently, cells were stimulated with 3 μg/ml OKT3 for 5 min and lysed. Cell extracts were analyzed by immunoblotting with antibodies recognizing the proteins indicated. The results are representative of three independent experiments. (**D**) Serum-starved Jurkat T cells were incubated with or without 1 μg/ml sec49K-His for 30 min and stimulated with 3 μg/ml OKT3. Cells were lysed at different time points after stimulation as indicated. Cell extracts were analyzed by immunoblotting with antibodies recognizing the proteins indicated. The results are representative of three independent experiments. (**E**) Calcium flux was measured in Jurkat T cells with Fluo-4 as a calcium sensor after 30 min of incubation with 1 μg/ml sec49K-His and stimulation with 3 μg/ml OKT3 for 0-30 min. Subsequently, the cells were lysed. Cell extracts were analyzed by immunoblotting with antibodies recognizing the proteins indicated. The mean fluorescence intensity (MFI) of mock treated cells was set as 1. The error bars show the standard deviation of the mean of three independent experiments. (**F**) Serum-starved Jurkat T cells were incubated with 1 μg/ml sec49K-His for 30 min and stimulated with 1 μg/ml OKT3 for 90 min. CD69 upregulation was determined by flow cytometry. The mean fluorescence intensity (MFI) of upregulated CD69 after TCR stimulation of mock treated cells was set as 1. The error bars show the standard deviation of the mean of four independent experiments (*P<1×10^−7^).

We previously demonstrated that sec49K impairs phosphorylation of ZAP70 and the extracellular signal-regulated kinases 1 and 2 (ERK1/2) after TCR stimulation by CD3 crosslinking in Jurkat T cells (Windheim et al., 2013). However, the most upstream substrate of CD45 in TCR signaling is the tyrosine kinase Lck in which CD45 removes the inhibitory phosphorylation at Y505 (Sieh et al., 1993). Therefore, we tested whether sec49K-His influenced the phosphorylation of Y505 Lck. Indeed, sec49K-His increased the basal levels of Y505-Lck phosphorylation in Jurkat T cells (Fig. 1B) and inhibited the dephosphorylation of pY505 Lck upon TCR stimulation with an anti-CD3ε antibody (OKT3) in a dose-dependent manner (Fig. 1C, D). The inhibition of pY505 Lck dephosphorylation correlated with the inhibition of ERK1/2 phosphorylation and activation (Fig. 1C, D). Another important signal for T cell activation is the increase of intracellular calcium (Trebak & Kinet, 2019). However, we did not observe an influence of sec49K-His on the calcium flux induced by TCR stimulation, although we detected an effect on the dephosphorylation of Lck and the phosphorylation of ERK1/2 in the sec49K-treated cells at 5-30 min after stimulation (Fig. 1E). Even when we performed experiments with continuous measurement of the intracellular calcium concentration (Vines et al., 2010) using lower anti-CD3ε antibody (OKT3) concentrations for TCR stimulation (0.1-3 μg/ml) and shorter incubation times (0-5 min), we did not detect a significant influence of sec49K-His on the calcium flux (data not shown).

We previously reported a negative effect of sec49K on the upregulation of the early activation marker CD69 after TCR stimulation (Windheim et al., 2013). In accord with this result, recombinant sec49K-His reduced the upregulation of CD69 in Jurkat T cells after TCR stimulation significantly by about 40% (Fig. 1F). In summary, we demonstrate here that the recombinant sec49K-His expressed in insect cells strongly reduced both Lck dephosphorylation and ERK phosphorylation after TCR stimulation and, therefore, appears to exhibit the same properties as native sec49K. Since protein glycosylation in insect and mammalian cells differs significantly (van Oers et al., 2015), we conclude that specific glycan structures are not critical for the binding of sec49K to CD45 and the sec49K-mediated inhibition of TCR signaling.

### E3/49K binds to the domain d3 present in all CD45 isoforms

The extracellular part of CD45 is divided into several distinct domains. It consists of an N-terminal mucin-like part with multiple sites of O-glycosylation comprising a stretch of 41 residues present in all CD45 isoforms and the variable regions A, B and C present only in some CD45 isoforms, and the membrane-proximal domains d1-d4, which are present in all CD45 isoforms (Fig. 2A, top) (Chang et al., 2016). The domains d1-d4 were formerly classified as a cysteine-rich domain and three fibronectin type 3 domains (FN3) (Rheinlander et al., 2018). However, recent structural analysis revealed an FN3 fold in all four domains, albeit with a degenerate topology in d1 and d2 (Chang et al., 2016). To determine which domains of CD45 interact with E3/49K, we replaced the intracellular phosphatase domains D1 and D2 of CD45RO (lacking regions A, B and C) and CD45RABC with GFP to create RO ECD-GFP and RABC ECD-GFP constructs. Subsequently, we deleted one or several of the extracellular domains of CD45 (Fig. 2A, Table 1). To test whether the variable regions A, B and C of CD45 are involved in binding to 49K, we transfected constructs encoding wild-type CD45RO and CD45RABC (Fig. 2B, C) or CD45RO ECD-GFP and CD45RABC ECD-GFP (Fig. 2D, E) into 293T cells. Sec49K-His binding was then investigated by flow cytometry (Fig. 2B-E). No significant difference in sec49K-His binding was detectable between wild-type CD45RABC and CD45RO (Fig. 2B, C) and between the CD45RABC ECD-GFP and CD45RO ECD-GFP proteins (Fig. 2D, E). Thus, the lack of the variable domains A, B and C had no significant effect on the binding of sec49K-His. Consequently, sec49K binds to a domain present in all CD45 isoforms, as previously suggested by antibody blocking studies (Windheim et al., 2013).

**Figure 2.**
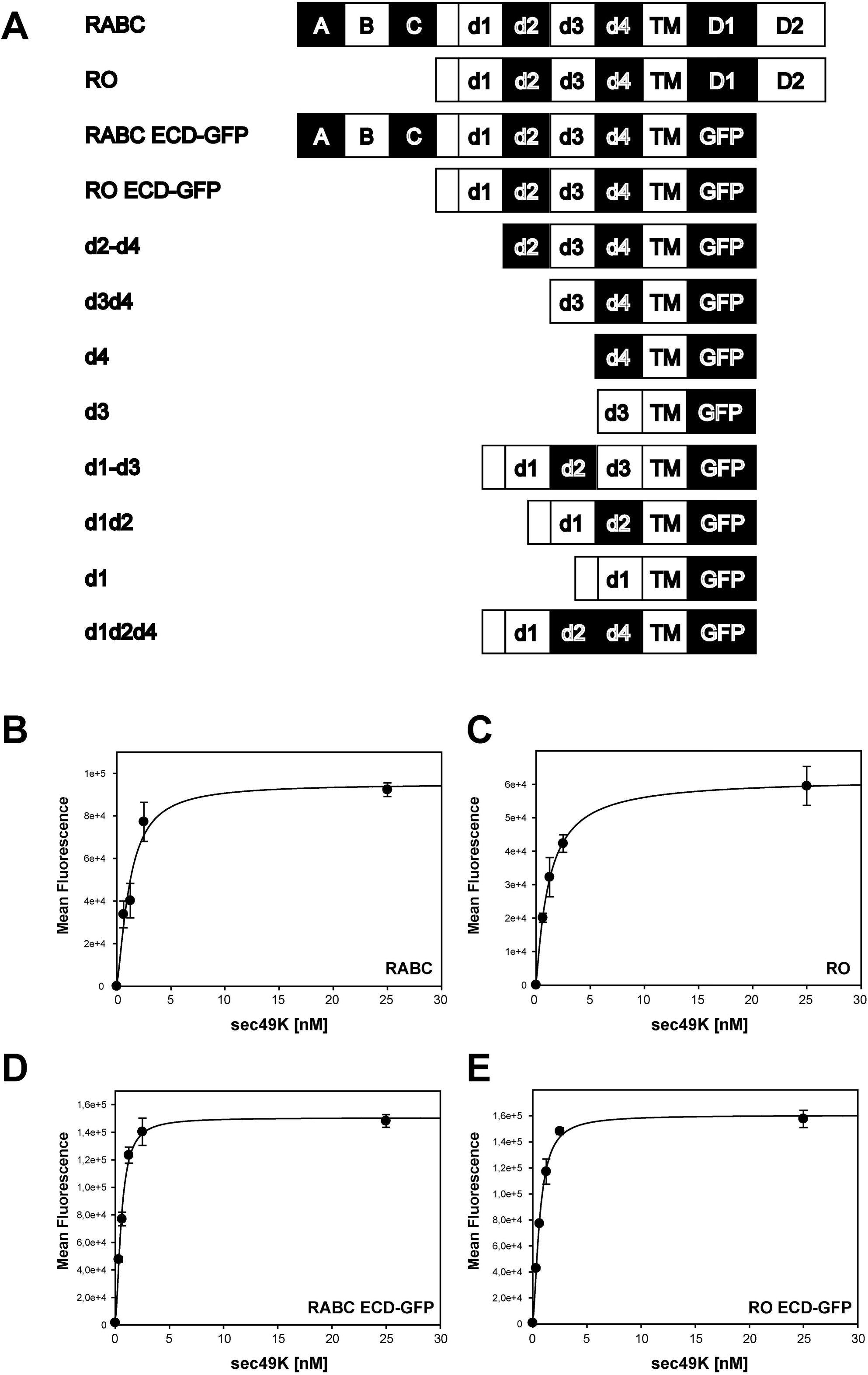
Sec49K binding to CD45 is not dependent on the variable regions A, B and C of CD45. (**A**) Schematic representation of the full-length human CD45RABC and RO and the constructs with the intracellular phosphatase domains replaced by GFP with or without deletions of one or several domains used in this study (see also Table 1). (**B-E**) 293T cells were transfected with constructs encoding full-length CD45RABC (**B**) and CD45RO (**C**) or CD45RABC ECD-GFP (**D**) and CD45RO ECD-GFP (**E**) with the phosphatase domains replaced by GFP. After 48 h, cells were detached and binding to sec49K-His was analyzed with an anti-49K ECD rabbit antiserum (R48) by flow cytometry. The expression levels of full-length CD45RABC and CD45RO or CD45RO ECD-GFP and CD45RABC ECD-GFP, respectively, were similar. The expression of full-length CD45 was measured with an anti-CD45 antibody (MEM28, data not shown). The expression of the GFP-tagged proteins was determined by GFP fluorescence. The results are representative of at least three independent experiments. The error bars indicate the standard deviation of the mean of two or three measurements. The mean EC50 values were 1.26 (+/- 0.51) nM for CD45RABC (**B**, n=4) and 1.35 (+/- 0.76) nM for CD45RO (**C**, n=4). The mean EC50 values were 0.55 (+/- 0.01) nM for CD45RABC ECD-GFP (**D**, n**=**3) and 0.54 (+/- 0.1) nM for CD45RO ECD-GFP (**E,** n=11).

**Table 1:** Overview of CD45 ECD deletion mutants and sec49K binding.

| **CD45 ECD mutant** | **CD45 Sequence\***<br>**Amino acid residues** | **Cell surface**<br>**binding\*\*** | **Ni NTA**<br>**Pulldown** |
| --- | --- | --- | --- |
| RABC | 25-630 | + | + |
| RO | 25-31, 193-630 | + | + |
| d2-d4 | 293-630 | - | + |
| d3d4 | 385-630 | - | + |
| d4 | 479-630 | - | (+)*** |
| d3 | 385-483, 569-630 | - | (+)*** |
| d1 | 25-31, 193-301, 569-630 | - | - |
| d1d2 | 25-31, 193-393, 569-630 | - | - |
| d1-d3 | 25-31, 193-483, 569-630 | + | + |
| d1d2d4 | 25-31, 193-393, 474-630 | - | - |

\* Receptor-type tyrosine-protein phosphatase C isoform 1 precursor NCBI Reference Sequence NP\_002829.2: Full-length (1304 residues), extracellular part (residues 1-577), transmembrane region (residues 578-600), tyrosine phosphatase domain 1 (residues 705-905), tyrosine phosphatase domain 2 (residues 1015-1221). The CD45 signal peptide (residues 1-24) was replaced by the signal peptide of CD4 (27 residues). The ECD deletion mutants contain in addition to the designated domains residues 569-630 encompassing the region between d4 and the transmembrane domain, the transmembrane domain and a short stretch of the intracellular part of CD45 (569-630) and may contain the N-terminal region between the signal peptide and d1 (25-31 and 193-225) of CD45RO.
\*\* determined by flow cytometry
\*\*\*nonspecific binding to the Ni-NTA beads possibly caused by misfolding

To determine the domain of CD45 sec49K binds to, we lysed transfected 293T cells expressing the CD45 ECD-GFP constructs (cf. Fig. 2A, Table 1). Subsequently, we incubated the cell extracts with recombinant sec49K-His, performed a pulldown with Ni-NTA agarose beads and determined whether the CD45 deletion mutants co-precipitated with sec49K-His. All proteins were expressed, albeit at different expression levels (Fig. 3A, B). The GFP-tagged ECDs of both, CD45 RABC and RO, were precipitated with sec49K-His with similar efficiency (Fig. 3C, left panel), demonstrating again that the variable domains A, B and C are not required for binding. In addition, the deletion mutants d2-d4, d1-d3 and d3d4 were pulled down with sec49K-His, whereas the CD45 deletion mutants d1 and d1d2 were not precipitated (Fig. 3C). The d3d4 mutant displayed some nonspecific binding to the beads. However, in the presence of sec49K-His a much stronger binding was observed demonstrating specific binding of d3d4 to sec49K-His (Fig. 3C, right panel). To confirm these findings, we mixed cell extracts containing the GFP-tagged CD45 ECD deletion mutants with extracts from cells expressing 49K full-length protein. Subsequently, the CD45 mutants were precipitated with a His-tagged nanobody recognizing GFP and Ni-NTA agarose beads and the co-precipitation of 49K was assessed (Fig. 3D). In accord with the previous results, 49K bound to the ECDs of CD45 RABC and RO, d2-d4, d1-d3 and d3d4, but not to d1 and d1d2 proteins (Fig. 3D). In summary, all deletion mutants containing d3 bound to 49K or sec49K whereas d1 and d2 were not able to bind. However, based on these data, a contribution of the domain d4 to sec49K binding could not be completely ruled out. Therefore, we also tested whether we could visualize binding of the single domains d3 and d4 to sec49K-His, but observed nonspecific binding to the Ni-NTA agarose beads in the absence of sec49K-His, possibly suggesting that these proteins are misfolded (Fig. 3E). Thus, it appears to be difficult to stably express the domains d3 and d4 individually, which has been observed before in attempts to express deletion mutants of rat CD45 (Symons et al., 1999). In an alternative approach to assess whether sec49K binds to the domain d4 of CD45, we deleted the domain d3 to express a d1d2d4 protein and compared that to the d1-d3 protein in the sec49K-His pulldown assay (Fig. 3F). Both proteins were expressed in 293T cells (Fig. 3B), but only the d1d2d3 protein bound to sec49K-His (Fig. 3F). Therefore, we conclude that only domain d3 of CD45 binds to 49K and domain d4 is not involved in binding. In addition, we tested whether we detect sec49K-His binding to the CD45 deletion mutants at the cell surface by flow cytometry (data not shown). However, sec49K-His binding at the cell surface could only be detected with the d1-d3 mutant in addition to RABC ECD-GFP and RO ECD-GFP (Table 1), suggesting that in the other sec49K-binding CD45 mutants identified in the Ni-NTA pulldown assay, d2-d4 and d3d4, the deletions impair the transport to the cell surface.

**Figure 3.**
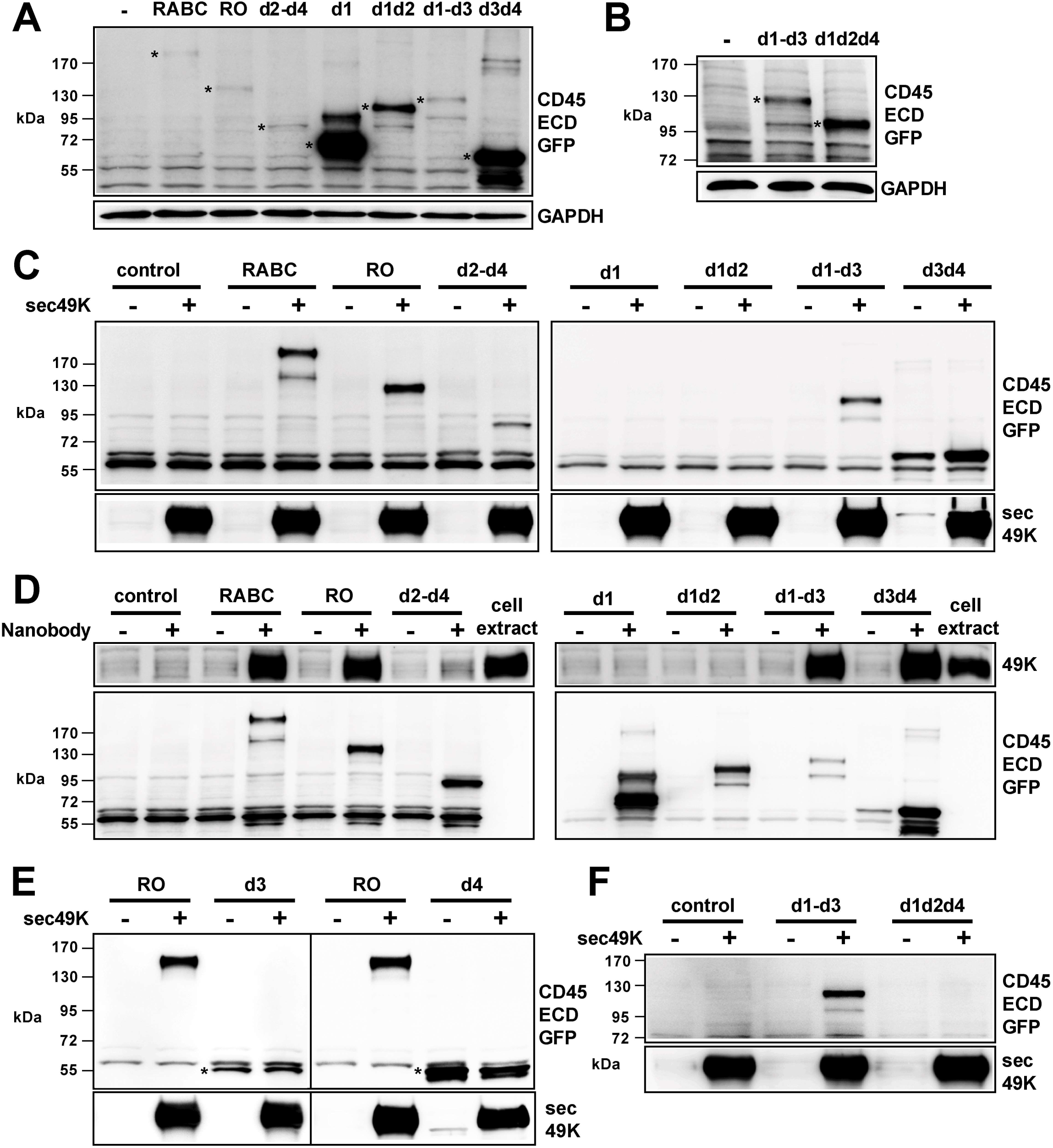
E3/49K binds to the domain d3 of CD45 present in all CD45 isoforms. (**A, B**) 293T cells were transfected with various CD45 ECD-GFP constructs. After 48 h, the cells were lysed and cell extracts were analyzed by immunoblotting with antibodies recognizing GFP to detect the CD45 ECD-GFP proteins and GAPDH as a loading control. Major bands representing the CD45 ECD-GFP proteins are marked with an asterisk (*). (**C, E, F**) Cell extracts (0.5 mg) were incubated with 1 μg sec49K-His. Proteins were precipitated with Ni-NTA agarose beads and were analyzed by immunoblotting with anti-GFP antibodies to detect the co-precipitated CD45 ECD mutants and with an anti-49K ECD rabbit antiserum (R48) to detect sec49K-His. The result is representative of at least three independent experiments. (**D**) Cell extracts (0.5 mg) from cells transfected with the CD45 ECD-GFP constructs were mixed with cell extracts (0.5 mg) from cells transfected with pSG5-49K expressing 49K and 2 μg of a His-tagged nanobody recognizing GFP. Proteins were precipitated with Ni-NTA agarose beads and were analyzed by immunoblotting with anti-GFP antibodies to detect the precipitated CD45 ECD mutants and with an anti-49K ECD rabbit antiserum (R48) to detect the co-precipitated 49K. Cell extract (20 μg) from cells transfected with pSG5-49K was analyzed as a control (cell extract). The result is representative of at least three independent experiments. (**E**) The bands corresponding to GFP-tagged d3 and d4 are marked with an asterisk (*).

### E3/49K contains at least two binding sites for CD45

Having identified the domain d3 of CD45 as the binding site for E3/49K, we next addressed the question of which domains of 49K are involved in the binding of CD45. HAdV-D64 49K consists of the extracellular domains R1 (residues 24-102), R2 (residues 119-197), R3 (residues 268-342), a longer stretch of about 60 amino acids (D, residues 203-260) linking R2 und R3 predicted to be disordered (Barik et al., 2020), the transmembrane region (TM, residues 386-409) and a short cytoplasmic tail (CT, residues 410-428, Fig. 4A, top) (Deryckere & Burgert, 1996). The N-terminal ectodomain is cleaved at the cell surface at a site between the R3 domain and the transmembrane region resulting in the shedding of the ECD (Windheim et al., 2016; Windheim et al., 2013). To determine which domains are involved in CD45 binding, we deleted one or two domains and transfected 293T cells with the constructs encoding the deletion mutants (Fig. 4A, Table 2). All proteins could be easily detected in the cell extracts (Fig. 4B, upper panel) and at the cell surface by flow cytometry (data not shown). Moreover, all these proteins were proteolytically processed, since the C-terminal fragments were detected in the cell extracts (Fig. 4B, middle panel) and the secreted N-terminal ECDs were efficiently shed into the supernatants of the transfected cells (Fig. 4C). To determine whether these 49K mutants bind CD45, we mixed the cell extracts (Fig. 4B) with a cell extract from 293T cells transfected with a construct encoding CD45RO ECD-GFP (cf. Fig. 3A). Subsequently, we precipitated the GFP-tagged ECD of CD45RO with a His-tagged nanobody recognizing GFP and Ni-NTA agarose beads. Co-precipitated 49K proteins were detected with the antibody recognizing the C-terminus, i.e. that the recognition is independent of the deletions in the N-terminal region (Fig. 4D, E). Specific binding to CD45RO ECD-GFP was demonstrated for 49K, R1R2, R2R3, R1R3 (-/+D, disordered region) and to a lesser extent to R1 (Fig. 4D, E). The single domain proteins R2 and R3 bound nonspecifically to the beads (without the nanobody), indicating possibly misfolding, and the addition of the nanobody did not increase the amounts of precipitated R2 or R3 (Fig 4E, left panel). However, although R2 and R3 alone did not interact with CD45, at least one of these domains must bind to CD45, since the R2R3 mutant co-precipitated with CD45RO ECD-GFP (Fig. 4D, upper panel). The deletion of the disordered region (-D) did not influence the binding of the R1R3 protein to CD45RO ECD (Fig. 4E, right panel). Furthermore, the deletion of the disordered region had no significant effect on the binding of sec49K to Jurkat T cells (data not shown). Thus, the disordered region appears not to be directly involved in CD45 binding. Taken together, we conclude that 49K contains at least two binding sites, one in the R1 domain and another one in R2 or R3.

**Figure 4.**
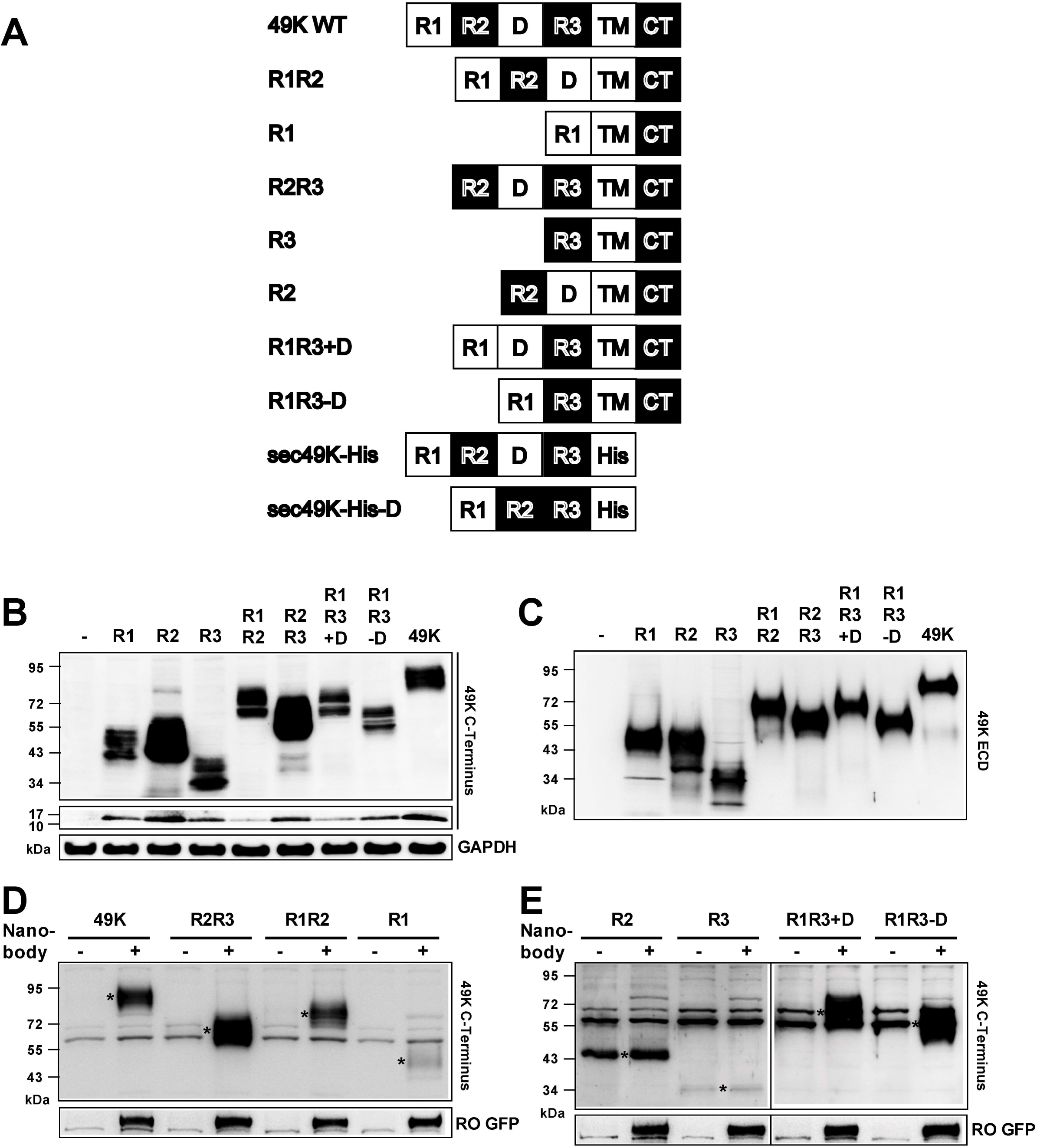
E3/49K contains at least two binding sites for CD45. (**A**) Schematic representation of full-length HAdV-D64 49K (49K WT) with three extracellular domains R1, R2 and R3, the disordered region (D), the transmembrane domain (TM) and the cytoplasmic tail (CT) and the constructs with deletions of one or several domains used in this study (see also Table 2). In the constructs encoding sec49K-His and sec49K-His-D a C-terminal 6xHis-tag (His) was added. (**B-E**) 293T cells were transfected with constructs encoding full-length 49K or deletion mutants. (**B**) After 48 h, cells were lysed. Cell extracts were analyzed by immunoblotting with an anti-49K C-terminus rabbit antiserum (25050), thereby detecting the 49K full-length protein and the deletion mutants (upper panel) and the cleaved cytoplasmic tail (middle panel). GAPDH was used as a loading control. (**C**) Supernatants were analyzed by immunoblotting with an anti-49K ECD rabbit antiserum (R518). To adjust for the different expression levels of the deletion mutants the following volumes of the supernatants were analyzed: 20 μl R1 and R3, 4 μl R1R3 (+/-D), 2 μl R2, R1R2, R2R3 and 49K. (**D, E**) Cell extracts (0.25 mg) from cells transfected with the CD45RO ECD-GFP construct were mixed with cell extracts (0.25 mg) from cells transfected with pSG5-49K full-length (49K) or the deletion mutants. Only 50 μg cell extract from cells expressing R2 was used because of the high expression level of R2. The CD45RO ECD-GFP protein was precipitated with 2 μg of a His-tagged nanobody recognizing GFP bound to Ni-NTA agarose beads. The precipitated proteins were analyzed by immunoblotting with anti-GFP antibodies to detect the CD45RO ECD-GFP protein and with an anti-49K C-terminus rabbit antiserum (25050) to detect 49K full-length and the deletion mutants. Co-precipitated 49K and deletion mutants are marked with an asterisk (*).The result is representative of at least five independent experiments.

**Table 2:** Overview of 49K deletion mutants.

| **49K or deletion mutant** | **Amino acid residues** |
| --- | --- |
| 49K full-length* | 1-428 |
| R1R2** | 1-267, 343-428 |
| R1** | 1-118, 343-428 |
| R2R3** | 1-23, 103-428 |
| R3** | 1-23, 261-428 |
| R2** | 1-23, 103-267, 343-428 |
| R1R3+D** | 1-118, 198-428 |
| R1R3-D*** | 1-118, 261-428 |
| sec49K-His | 1-353 |
| sec49K-His-D*** | 1-202, 261-428 |

\*The 49K sequence contains the following regions:
Signal peptide (residues 1-23), R1 (residues 24-102), R2 (residues 119-197), disordered region (residues 203-260), R3 (residues 268-342), transmembrane region (residues 386-409) and cytoplasmic tail (residues 410-428).
\*\*The 49K deletion mutants contain in addition to the designated domains the signal peptide (residues 1-23) and the region between R3 and the C-terminus (residues 343-428) and they may contain the regions between R1 and R2 (residues 103-118) or R2 and R3 (residues 198-267).
\*\*\*The region between residues 203 and 260 was deleted in 49K mutants lacking the disordered region (-D).

### E3/49K domains R1 and R2 enforce dimerization of CD45

To determine how many CD45 binding sites are present in E3/49K, we performed crosslinking experiments using two crosslinkers with spacer arms of different lengths, BS^2^G-d_0_ (“2”) with a spacer arm of 7.7 Å and BS^3^-d_0_ (“3”) with a spacer arm of 11.4 Å. Initially, in crosslinking experiments in Jurkat T cells and 293T cells overexpressing CD45RO we detected substantial CD45 crosslinking even in the absence of sec49K (data not shown). However, in 293T cells overexpressing CD45RO ECD-GFP only very low levels of crosslinked CD45 potentially representing CD45 dimers were detected (Fig. 5A, upper panel; -sec49K, +2 and +3). Consequently, we continued with this system to investigate sec49K-mediated CD45 crosslinking.

**Fig. 5.**
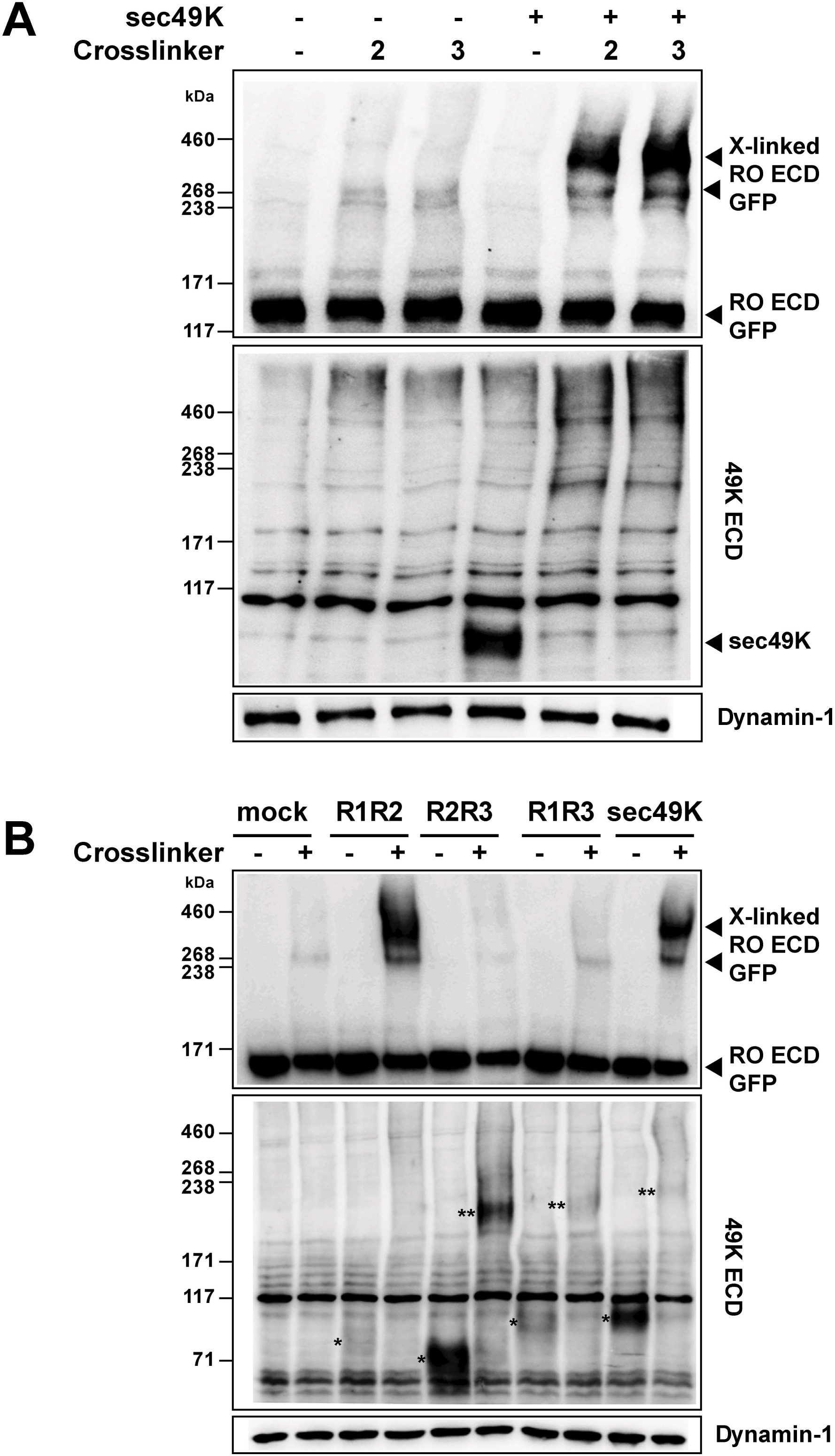
E3/49K domains R1 and R2 enforce dimerization of CD45. (**A**) 293T cells were transfected with a construct encoding CD45RO ECD-GFP and detached after 48 h. Cells were incubated with a supernatant from pSG5-49K transfected cells containing sec49K. Crosslinking of CD45RO and 49K was performed with the crosslinker BS^2^G-d_0_ (“2”, spacer arm of 7.7 Å) and with the crosslinker BS^3^-d_0_ (“3”, spacer arm of 11.4 Å). Subsequently, the cells were lysed and the cell extracts were analyzed by immunoblotting with antibodies recognizing GFP (upper panel), the 49K ECD (R518, middle panel) and dynamin-1 as a loading control. Bands representing CD45RO ECD-GFP and the crosslinked (“X-linked”) CD45RO ECD-GFP complexes are marked with arrowheads. The result is representative of three independent experiments. (**B**) 293T cells were transfected with a construct encoding CD45RO ECD-GFP and detached after 48 h. Cells were incubated with supernatants from cells transfected with pSG5 constructs encoding 49K full-length (sec49K) or deletion mutants lacking the domains R3 (R1R2), R1 (R2R3) or R2 (R1R3). Crosslinking of CD45RO ECD-GFP was performed with the crosslinker BS^3^-d_0_. Subsequently, the cells were lysed and cell extracts were analyzed by immunoblotting with antibodies recognizing GFP (upper panel), the 49K ECD (R518, middle panel) and dynamin-1 as a loading control. Bands representing CD45RO ECD-GFP and the crosslinked (“X-linked”) CD45RO ECD-GFP complexes are marked with arrowheads. Bands representing R1R2, R2R3, R1R3, 49K proteins recognized by the anti-49K ECD rabbit antiserum (R518) are marked with an asterisk (*) and their crosslinked products are marked with two asterisks (**). The result is representative of three independent experiments.

Thus, sec49K-containing supernatants were incubated with 293T cells expressing CD45RO ECD-GFP and crosslinking was performed. Subsequently, cells were lysed and the crosslinked complexes were analyzed (Fig. 5A). In the absence of crosslinkers only the CD45RO ECD-GFP is detected with an apparent molecular weight of 130-150 kDa (Fig. 5A, upper panel). After addition of the crosslinkers, a weak band appeared with an apparent molecular weight of approximately 270 kDa in the absence of sec49K, which may represent CD45 dimers (Fig. 5A, upper panel; -sec49K, +2 and +3). Strikingly, in the presence of sec49K strong bands of crosslinked CD45-containing complexes with an apparent molecular weight of 340-460 kDa were detected with both crosslinkers (Fig. 5A, upper panel; +sec49K, +2 and +3). The apparent molecular weight of these complexes is consistent with a dimer of CD45 crosslinked to one sec49K protein. However, although the sec49K protein was detected bound to CD45RO ECD-GFP before crosslinking (Fig. 5A, middle panel; +sec49K, -crosslinker), there was no signal detected with an antibody directed against the N-terminal ECD of 49K at the same apparent molecular weight where the anti-GFP antibody detected the crosslinked complex (Fig. 5A; +sec49K, +2 and +3; compare upper and middle panel). Thus, either sec49K is not present in the complex or it is not recognized in the complex by the antibody because the epitopes are not accessible or destroyed after crosslinking. In favor of the latter, sec49K was clearly recognized prior to crosslinking (Fig. 5A, middle panel; +sec49K, -crosslinker), but not after crosslinking (Fig. 5A, middle panel; +sec49K, +2 and +3) suggesting that it was indeed crosslinked to CD45 RO ECD-GFP. Taken together, sec49K induced a complex that based on its apparent molecular weight could be a 49K-containing dimer of CD45 or a 49K-less trimer of CD45.

To distinguish between these alternatives and to determine whether R2 or R3 or both bind to CD45, we performed crosslinking experiments with supernatants containing mutants of 49K with two domains, i.e. R1R2, R2R3 and R1R3. Full-length sec49K induced the formation of a CD45-containing complex as shown before (Fig. 5B, upper panel; sec49K, +crosslinker). In the presence of R2R3 and R1R3 such a crosslinked complex was not detected (Fig. 5B, upper panel; R2R3 and R1R3, +crosslinker). However, both proteins bound to CD45RO ECD-GFP (Fig. 5B, middle panel; R2R3 and R1R3, -crosslinker (*)) and also some R2R3- and R1R3- containing aggregates were detected with an antibody recognizing the N-terminal ECD of 49K (Fig. 5B, middle panel; R2R3 and R1R3, +crosslinker (**)). But the apparent molecular weight of these aggregates was below 238 kDa and they appeared not to contain CD45RO ECD-GFP, because they were not detected with the antibody recognizing GFP (Fig. 5B, upper panel; R2R3 and R1R3, +crosslinker). Strikingly, the R1R2 protein was able to induce the formation of a CD45-containing complex of an apparent molecular weight similar to that detected with the full-length sec49K protein (Fig. 5B, upper panel; R1R2, +crosslinker). However, as seen before, this high molecular weight complex could not be detected with an antibody directed against the N-terminal ECD of 49K (Fig. 5B, middle panel; R1R2, +crosslinker). Nonetheless, these results suggested that the full-length 49K and the R1R2 proteins contain two CD45 binding sites and trigger CD45 dimerization. Thus, the N-terminal domains R1 and R2 both contain a binding site for CD45, whereas the membrane-proximal domain R3 does not bind CD45.

To determine whether R1 and R2 bind to the same site on CD45 or to two distinct binding sites, we performed competition experiments with Jurkat T cells expressing CD45 and supernatants containing 49K deletion mutants with only two domains. These were used to compete with supernatant containing sec49K-His, which could be detected with an anti-His antibody. If there were two different binding sites in CD45, a competitor that binds to only one site would not be able to efficiently block the binding of sec49K-His. However, if there were only one binding site in CD45, all 49K mutants, including the ones with only one CD45 binding domain, would be able to efficiently block sec49K-His binding. Evidently, all the 49K deletion mutants, R1R2, R2R3 and R1R3, were able to compete efficiently with sec49K-His for the binding to Jurkat T cells (Fig. 6A). Thus, we conclude that R1 and R2 bind to the same binding site in CD45.

**Fig. 6.**
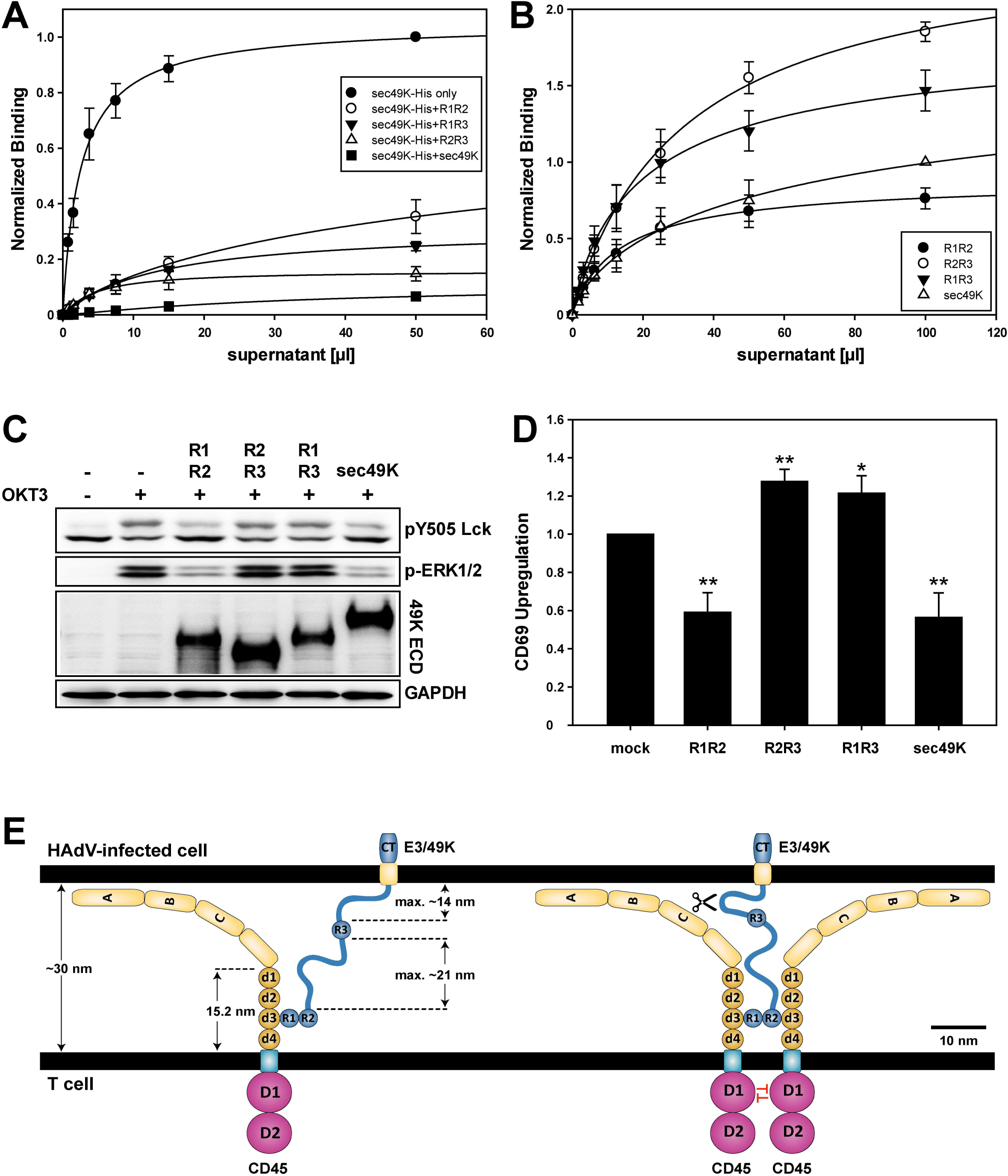
Sec49K-mediated inhibition of TCR signaling requires two CD45-binding domains. (**A**) Jurkat T cells were incubated for 45 min with 100 μl supernatant from 293T cells transfected with pSG5 constructs encoding 49K full-length (sec49K) or deletion mutants lacking R3 (R1R2), R2 (R1R3) or R1 (R2R3). The binding capacity of the supernatants was determined before to ensure that 100 μl supernatant are sufficient for at least 75% saturation of the sec49K binding sites on Jurkat T cells (5×10^5^ cells). Subsequently, different volumes of supernatant from 293T cells transfected with a pSG5 construct encoding sec49K-His (see Fig. 4A) were added and incubated for 45 min. Sec49K-His was detected with an anti-His antibody by flow cytometry. The measured mean fluorescence intensity (MFI) using 50 μl His-tagged sec49K supernatant was set as 1. The error bars show the standard deviation of the mean of three independent experiments. (**B**) Jurkat T cells were incubated with supernatants from cells transfected with pSG5 constructs encoding 49K full-length (sec49K) or deletion mutants lacking the domains R3 (R1R2), R2 (R1R3) or R1 (R2R3). Binding was analyzed by flow cytometry with the rat monoclonal antibody 4D1, which recognizes an epitope in the disordered region between R2 and R3. The mean fluorescence intensity (MFI) using 100 μl 49K supernatant was set as 1. The error bars show the standard deviation of the mean of five independent experiments. (**C**) Serum-starved Jurkat T cells were incubated for 30 min with supernatants from 293T cells transfected with pSG5 constructs encoding 49K full-length (sec49K) or deletion mutants lacking the domains R3 (R1R2), R2 (R1R3) or R1 (R2R3). Subsequently, cells were stimulated with 3 μg/ml OKT3 for 5 min and lysed. Cell extracts were analyzed by immunoblotting with the antibodies recognizing the proteins indicated. The results are representative of three independent experiments. (**D**) Serum-starved Jurkat T cells were incubated for 30 min with supernatants from 293T cells transfected with pSG5 constructs encoding 49K full-length (sec49K) or deletion mutants lacking the domains R3 (R1R2), R2 (R1R3) or R1 (R2R3) (Fig. 4C). The binding capacity of the supernatants was determined before to ensure that 100 μl supernatant are sufficient for at least 75% saturation of the sec49K binding sites on Jurkat T cells (5×10^5^ cells). Subsequently, the cells were stimulated with 1 μg/ml OKT3 for 90 min. CD69 upregulation was determined by flow cytometry. The mean fluorescence intensity (MFI) of upregulated CD69 after TCR stimulation of mock treated cells was set as 1. The error bars show the standard deviation of the mean of four independent experiments (*P<0.005, (**P<0.0005). (**E**) Model of the E3/49K-mediated enforced dimerization of CD45 and the inhibition of phosphatase activity. We previously demonstrated that E3/49K binds CD45 as soluble sec49K after shedding and inhibits T cell signaling and function. Here we show that E3/49K binds to the domain d3 of CD45 using the N-terminal domains R1 and R2. Calculating the relevant dimensions between CD45 d3 and E3/49K R1 and R2 suggests that E3/49K may additionally bind CD45 as a membrane-anchored full-length protein, as shown here, as well as in solution (after shedding) as sec49K. For the membrane-anchored protein the disordered region between R2 and R3 appears to be important to bridge the distance to the binding site in the domain d3 of CD45. The length of the domains d1-d4 of CD45 is 15.2 nm and the mucin-like region of the CD45 isoform RO contains 41 residues and is ∼ 8 nm long resulting in an effective height of ∼22 nm for RO (Chang et al., 2016). The regions A, B and C comprise 66, 47 and 48 residues, respectively. If a length of 0.2 nm per residue similar to RO is assumed, this results in a length of 13.2 (A), 9.4 (B) and 9.6 nm (C). If a contour length of 0.35 nm per residue is assumed for the disordered region of 49K, this would result in a length of ∼21 nm for the stretch of about 60 residues between R2 and R3 in an extended conformation. Accordingly, the region of 49K between the transmembrane domain and the R3 domain comprising about 40 residues would be ∼14 nm long. This region contains the proteolytic cleavage site (indicated by a pair of scissors, right). Thus, E3/49K appears to be designed as a “molecular fishing rod” using the elongated disordered regions as a “fishing line” and the CD45-binding domains R1 and R2 as baits to “catch” and dimerize CD45. The dimerization of CD45 may sterically block the access of substrates to the active sites of the intracellular phosphatase domains (D1) resulting in the inhibition of TCR signaling and T cell activation.

This result would suggest that proteins with one CD45 binding domain would theoretically require two times the molar amount of protein to saturate all CD45 binding sites on the cell surface of Jurkat cells compared to proteins with two binding sites, if potential differences in affinity or avidity were considered not significant. We tested this idea with a rat monoclonal antibody (4D1) that recognizes an epitope in the disordered region between R2 and R3 and, therefore, binds all 49K mutants containing this region with the same affinity. As anticipated, we found that mutants with two CD45 binding sites, R1R2 and 49K, exhibited indeed a lower fluorescence signal in flow cytometry at concentrations close to saturation than mutants with one CD45 binding site, R2R3 and R1R3 (Fig. 6B). Thus, this result further supports the idea that R1 and R2 are the two CD45 binding domains of 49K and that they bind to the same binding site on CD45, which is located in the domain d3 as shown before.

### Sec49K-mediated inhibition of TCR signaling requires two CD45-binding domains

The presented results point to a potential mechanism of the 49K-mediated inhibition of TCR signaling (Fig. 1). The two CD45-binding domains of 49K could enforce dimerization of CD45 in a way that imposes an orientation on the intracellular phosphatase domains that inhibits substrate access as suggested previously as a regulatory mechanism of CD45 activity (Desai et al., 1993; Takeda et al., 1992; Xu & Weiss, 2002). If this hypothesis is correct, only 49K variants with two CD45-binding domains, R1R2 and full-length 49K, should inhibit CD45 activity and TCR signaling, whereas deletion mutants with only one CD45 binding domain, R2R3 and R1R3, should not. Thus, we determined how the different 49K deletion mutants affect TCR signaling in Jurkat T cells. Strikingly, R1R2 and 49K reduced the dephosphorylation of the inhibitory pY505 Lck and the phosphorylation of ERK1/2 after TCR stimulation, whereas R2R3 and R1R3 had no significant effect (Fig. 6C).

In addition, we assessed the impact of the 49K deletion mutants on the upregulation of the early activation marker CD69 upon TCR stimulation. Full-length sec49K reduced CD69 upregulation by about 40% (Fig. 6D). This was similar to the reduction observed with recombinant sec49K-His (Fig. 1F). Strikingly, the R1R2 protein decreased CD69 upregulation to a similar extent, whereas both R2R3 and R1R3 were not able to block CD69 upregulation and even increased CD69 levels slightly (Fig. 6D). In summary, we show here that the immunomodulatory function of 49K requires two CD45 binding domains and propose that 49K inhibits CD45 by a mechanism of enforced dimerization.

## Discussion

In this study, we demonstrate that the binding of the HAdV-D64 protein E3/49K to the tyrosine phosphatase CD45 induces its dimerization through interactions of the R1 and R2 domains of E3/49K and the domain d3 of CD45. This sec49K-mediated dimerization impaired substrate dephosphorylation and TCR signaling. Thereby, sec49K increased the basal levels of Lck with the inhibitory pY505 phosphorylation and inhibited pY505 dephosphorylation upon TCR stimulation in Jurkat T cells. This resulted in a strongly reduced phosphorylation of the downstream target ERK1/2, but did not impair calcium flux. Using a readout that integrates TCR signaling from different signal transduction pathways, sec49K reduced the CD69 upregulation upon TCR stimulation in Jurkat T cells by about 40%. The levels of active Lck are considered an important threshold for TCR signaling with CD45 functioning as a “signaling gatekeeper”, suppressing antigen-independent and weak signals while providing a pool of active Lck for T cell activation (Courtney et al., 2019; Lovatt et al., 2006). The missing effect of sec49K on intracellular calcium levels may be explained by a lower threshold for calcium signaling than for ERK1/2 activation. In line with this hypothesis it was previously shown that even a single peptide:MHC complex can trigger suboptimal calcium flux and ∼10 complexes are sufficient for maximal calcium flux (Huang et al., 2013; Irvine et al., 2002; Purbhoo et al., 2004).

We propose that sec49K-induced dimerization impairs the phosphatase activity of CD45. Such a negative regulation of CD45 phosphatase activity by dimerization has been previously suggested (Desai et al., 1993; Majeti et al., 1998; Takeda et al., 2004; Takeda et al., 1992). However, the cytoplasmic phosphatase domains D1 and D2 as well as the ECDs of CD45RO and RABC were reported to be monomeric when expressed alone (Barr et al., 2009; Chang et al., 2016; Nam et al., 2005) suggesting an important role of the transmembrane region for CD45 dimerization. In support of this hypothesis, it was reported that the CD45-associated protein (CD45-AP) interacts with CD45 through its transmembrane region to impair CD45 dimerization (Cahir McFarland & Thomas, 1995; Kitamura et al., 1995). On the other hand, the extent of the dimerization was reported to differ between the CD45 isoforms, which would suggest an involvement of the ECD in this process (Dornan et al., 2002; Xu & Weiss, 2002). Moreover, *in vitro* binding studies suggested intermolecular and intramolecular interactions between the intracellular D1 and D2 domains (Felberg & Johnson, 1998, 2000). These seemingly contradictory results could be reconciled by adopting a “zipper” model of dimerization, in which multiple regions of the monomers interact in the dimer as proposed for RPTPα (Jiang et al., 2000). An inhibitory wedge was suggested to be important for the regulation of CD45 phosphatase activity by dimerization (Majeti et al., 1998; Majeti et al., 2000). However, the crystal structure of the D1 and D2 domains seems incompatible with the wedge model (Nam et al., 2005). Alternatively, a “head-to-toe” dimerization model has been proposed for RPTPγ/ζ with the D1 domain of one monomer interacting with the D2 domain of the second monomer (Barr et al., 2009), and, furthermore, another dimerization mode was recently shown for the leukocyte common antigen-related receptor protein tyrosine phosphatases (LAR-RPTPs), where two catalytic D1 domains interact via the “backside” of one monomer and the “frontside” of the other monomer blocking the substrate-binding pocket of the latter (Xie et al., 2020). Thus, although dimerization seems to regulate the activity of protein tyrosine phosphatases, it remains elusive whether a conserved inhibitory mechanism exists. Nevertheless, the E3/49K protein may have hijacked the physiological regulatory mechanism of dimerization to inhibit CD45 phosphatase activity.

Interestingly, other viruses also have been reported to target CD45, yet appear to use different strategies. The T-cell-tropic roseoloviruses downregulate CD45 transcripts and the MCMV m42 protein induces CD45 degradation in lysosomes in infected cells (Thiel et al., 2016; Whyte et al., 2021). The m42 protein is a tail-anchored membrane protein with a cytosolic N-terminal domain lacking a significant extracellular part (Thiel et al., 2016). The HCMV UL11 protein is a type I transmembrane protein with an extracellular part of ∼200 amino acids containing one RL11D domain with some homology to the E3/49K domains R1-R3 (Davison et al., 2003). The UL11 protein binds to an unknown region present in all CD45 isoforms, but with presumably only one CD45 binding site it would be unable to induce CD45 dimerization (Gabaev et al., 2011). However, a protein consisting of the extracellular part of the UL11 protein fused to the Fc fragment of human IgG reduced T cell signaling and proliferation (Gabaev et al., 2011) and induced the secretion of the anti-inflammatory cytokine IL-10 in CD4^+^ T cells (Osanyinlusi et al., 2022; Zischke et al., 2017). The IL-10 secretion induced by HCMV-infected epithelial cells in peripheral blood mononuclear cells and CD4^+^ T cells seemed to partially depend on the presence of the full-length transmembrane UL11 protein (Osanyinlusi et al., 2022; Zischke et al., 2017). However, the expression of the UL11 protein did not affect the activation of CD8^+^ T cells (Gabaev et al., 2014). Since the Fc fragment forms homodimers (Czajkowsky et al., 2012), the UL11 monomer is forced into a dimer with two CD45 binding sites in the recombinant UL11-Fc fusion protein. Thus, the UL11-Fc fusion protein unlike the monomeric full-length transmembrane protein may artificially induce CD45 dimerization similar to E3/49K to impair T cell function (Gabaev et al., 2011; Osanyinlusi et al., 2022; Zischke et al., 2017). It remains to be seen whether monomeric UL11 protein has indeed the same effects.

Using native sec49K (Windheim et al., 2013) and recombinant sec49K-His (this study), we clearly established that the secreted form of E3/49K is a negative regulator of immune cells. It is puzzling though why the virus does not simply encode a secreted protein, rather than expressing a transmembrane protein and generating a secreted version via shedding. This may suggest that the membrane-anchored full-length 49K functions similarly to sec49K. In addition, there appears to be no reason to assume that membrane-anchored 49K is not able to bind to and inhibit CD45 like the secreted sec49K unless steric constraints prevent this. A critical steric factor for CD45 binding of membrane-anchored 49K would be the size or more precisely the height of the CD45 ECD perpendicular to the plasma membrane, which would define the minimal size of the intercellular cleft. Membrane-anchored 49K would have to bridge the distance from the plasma membrane of the infected cell to the binding site in domain d3 of CD45 on the T cell or another leukocyte. That means, if the distance between the plasma membrane of the infected cell and the CD45-binding 49K domains R1 and R2 were smaller than the height of the N-terminal mucin-like regions and the domains d1 and d2 of CD45, binding of membrane-anchored 49K to the domain d3 of CD45 would be sterically impossible, the membrane-anchored 49K would be too “short” for binding (Fig. 6E). Thus, an assessment of the height of CD45 is important to evaluate this possibility.

Initially, the lengths of CD45RO and RABC ECDs were estimated by electron microscopy to be 28 nm and 51 nm, respectively (McCall et al., 1992). More recently, the length of the rigid linear d1-d4 domains was determined by crystallography to be 15.2 nm and a length of ∼22 nm for CD45RO was estimated by negative staining electron microscopy (Chang et al., 2016). The number of amino acids in the mucin-like regions of the CD45 isoforms ranges from 41 (CD45RO) to 202 residues (CD45RABC). For the mucin-like region of CD45RO a length of ∼8 nm was measured (Chang et al., 2016), i.e. approximately 0.2 nm per residue. With this estimate the 161 residues of the A, B and C regions in the CD45RABC isoform would be ∼32 nm long. However, apart from the length, the orientation of the protein between an upright and parallel position relative to the plasma membrane would determine the effective height, and this depends on a number of factors, such as the flexibility, the density of the proteins and the rotation around the anchoring point in the membrane (Junghans, Santos, et al., 2018). Surprisingly, using variable-angle total internal reflection fluorescence microscopy only a small difference in the effective height between CD45RO and CD45RABC was determined (Chang et al., 2016). Taken together, it was proposed that CD45RABC behaves like a 40 nm long rod that freely rotates around its anchoring point resulting in an effective height of ∼25 nm (Junghans, Hladilkova, et al., 2018). Interactions with neighboring proteins at the cell surface may increase the effective height by ∼5 nm to 30 nm (Junghans, Santos, et al., 2018) (Fig. 6E).

Considering these dimensions, membrane-anchored 49K would have to place its CD45- binding R1 and R2 domains over a distance of ∼22 nm for CD45RO and ∼30 nm for CD45RABC, reduced by ∼4 nm for the membrane-proximal domain d4, to enable binding to domain d3 of CD45. In this context, the part between R2 and R3 (residues 198-267) comprising the disordered region between residues 200 and 260, and the 43 residues between the transmembrane region and the R3 domain may be important. In an extended conformation, assuming a contour length of 0.35-0.4 nm per amino acid (Ainavarapu et al., 2007), the maximal length of these ∼100 residues would be 35-40 nm. The R3 domain with a length of ∼3.5 nm would further increase the distance between the transmembrane domain and the R1 and R2 domains of 49K and this would be clearly enough to allow binding to CD45 at the estimated intercellular cleft of ∼30 nm between the infected cell and the T cell (Fig. 6E). Intriguingly, the presence of an extended disordered region strongly supports the idea that the binding of membrane-anchored 49K to CD45 is indeed functionally important. In such a scenario, 49K may function as a “molecular fishing rod” to use the R3 domain and the disordered region as a “fishing line” and R1 and R2 as baits to “fish” for CD45 on the surface of T cells or other immune cells (Fig. 6E). In contrast, the disordered region would not be required for CD45 binding of shed sec49K. In accord with this assumption, the deletion of the disordered region in the secreted sec49K did not significantly affect CD45 binding suggesting that it is not directly involved in the interaction. In support of this idea, the disordered region is poorly conserved among HAdV-D 49K proteins (Blusch et al., 2002). Strikingly, the R3 domain is also less conserved than R1 and R2 and the homology between R1 and R2 (47%) is higher than that between R3 and R1 (36%) or R2 (29%) (Deryckere & Burgert, 1996; Martinez-Martin et al., 2016). Thus, the disordered region and possibly also the domain R3 may function mainly as a spacer or “fishing line” to enable binding of the domains R1 and R2 of the membrane-anchored 49K to the domain d3 of CD45 (Fig. 6E). Interestingly, the HCMV UL11 protein is predicted to contain also a disordered region (residues 139-230) (Barik et al., 2020) between the transmembrane domain and the RL11D domain (residues 40-107) (Gabaev et al., 2011), which may have a similar function. Taken together, a dual function of membrane-anchored and soluble 49K may be envisioned with the membrane-anchored 49K targeting immune cells that directly attack the infected cell, like cytotoxic CD8^+^ T cells or NK cells, while soluble sec49K would allow additionally the inhibition of CD45^+^ cells over a greater distance, e.g. CD4^+^ T cells or B cells.

Targeting CD45 phosphatase activity is regarded as a therapeutic strategy for the treatment of autoimmune diseases and organ transplantation (He et al., 2014; Stanford & Bottini, 2017). Moreover, CD45 could be targeted in cancer therapy, since it is often abnormally expressed in leukemia and lymphoma (Perron & Saragovi, 2018). Due to the specificity problems of small-molecule phosphatase inhibitors, biotherapeutics might be used as an alternative approach (Senis & Barr, 2018). Hence, recombinant sec49K could be applied as a biotherapeutic and small molecules or peptides may be developed that mimic E3/49K activity. Furthermore, CD45-binding E3/49K domains could be used for therapeutic strategies involving enforced phosphatase recruitment (Fernandes et al., 2020). Thus, further characterization of the E3/49K-mediated immunomodulation would be of great importance to evaluate the therapeutic potential of such strategies.

## Acknowledgments

We thank Christopher Tiedje (Hannover Medical School, Hannover, Germany) for the plasmid encoding the GFP nanobody. We thank David M. Rothstein (University of Pittsburgh, USA) for the plasmids encoding the CD45 isoforms RABC and RO. We thank Anneke Doerrie, Tim Dolgner and Marisa Bettels for excellent technical assistance. We thank Alexey Kotlyarov for helpful discussions and for critically reading the manuscript. Parts of this work were supported by the ‘Young Academy’ program of the Hannover Medical School.

## Materials and methods

All resources utilized in this study are listed below (Table 3) and referenced in the relevant methods section.

**Table 3.**
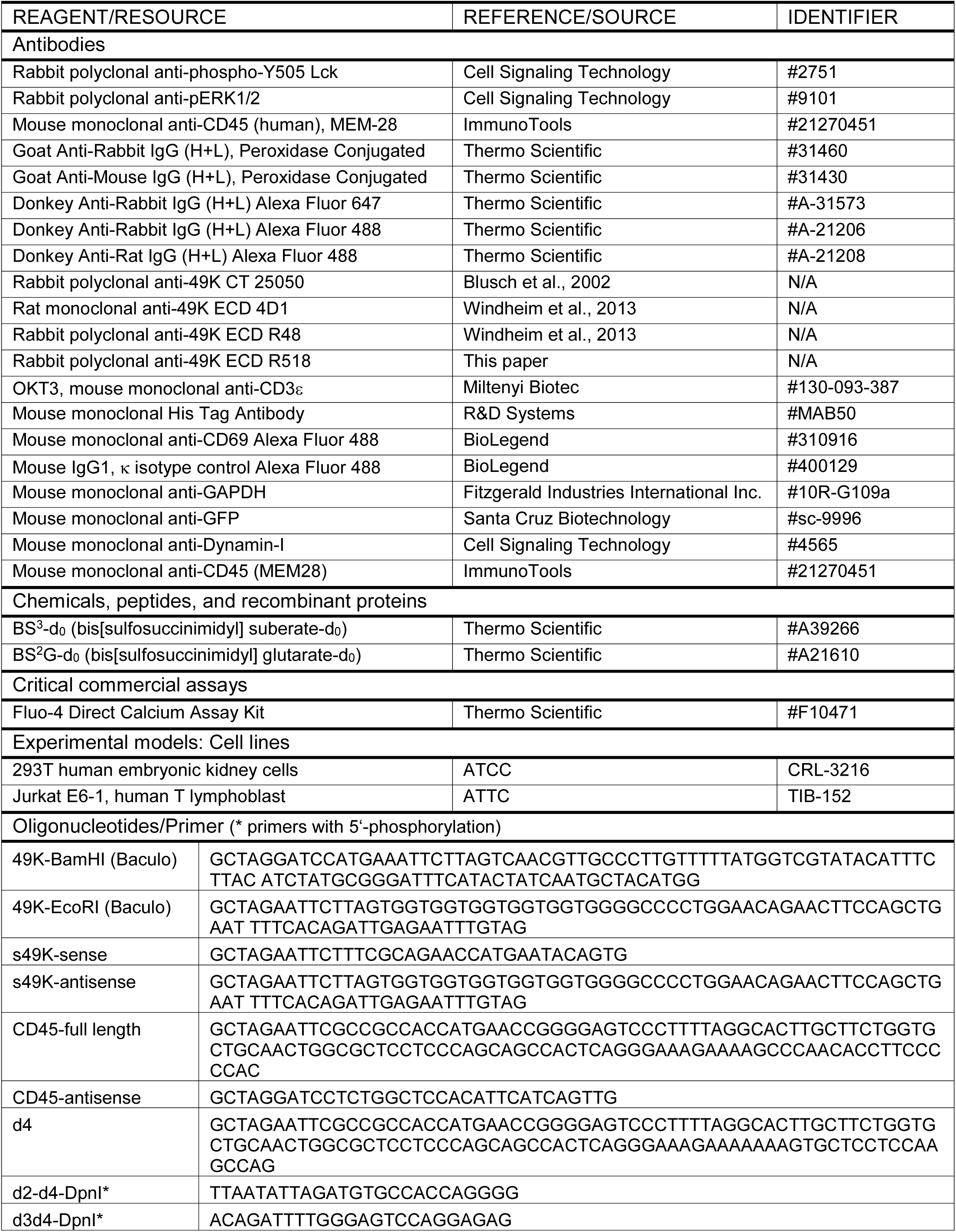
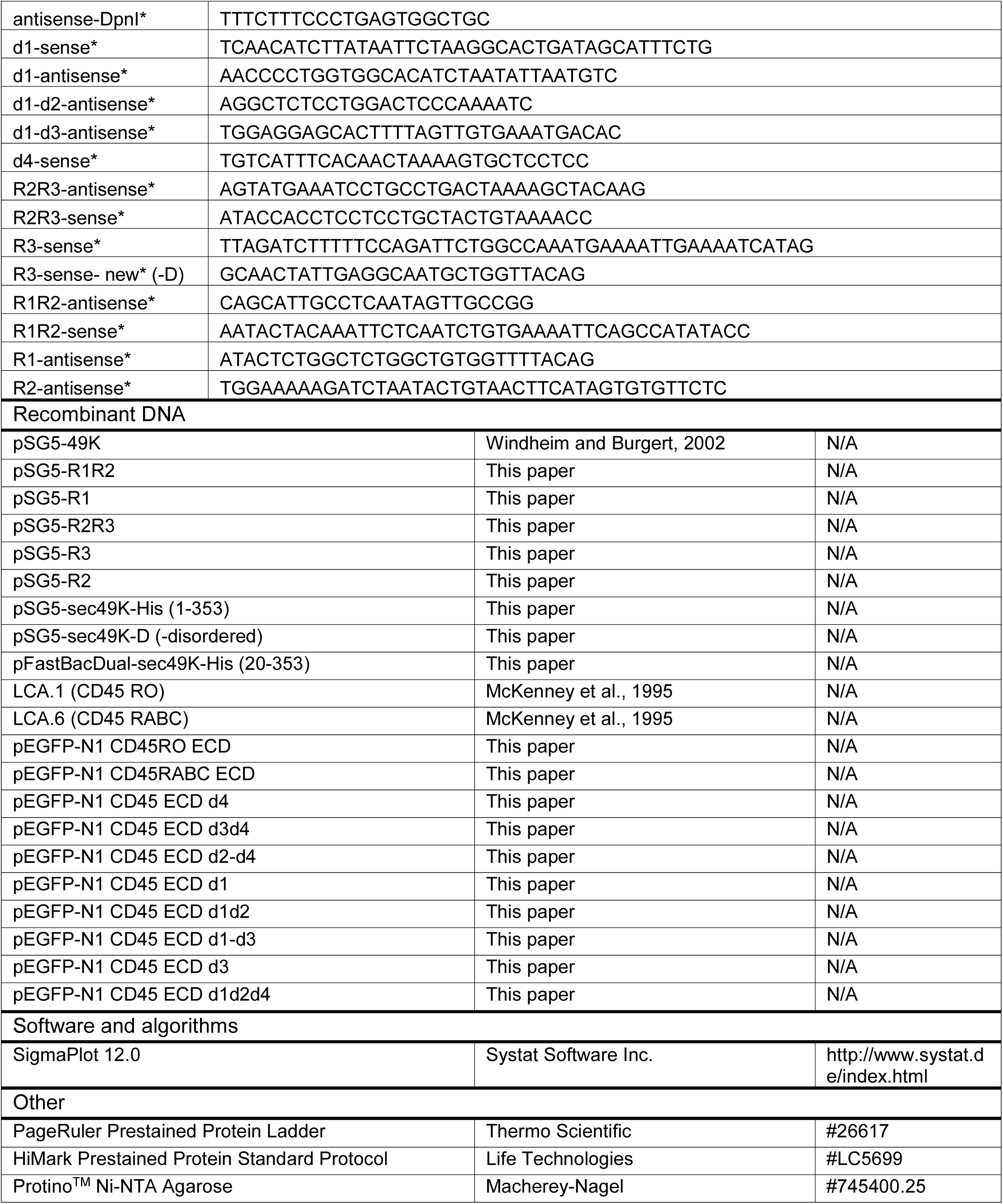
Key resources.

### Cell culture

293T cells (ATCC) were cultured in Dulbecco’s modified Eagle’s medium (DMEM) supplemented with 10% (v/v) fetal calf serum (FCS), 100 units/ml penicillin, 100 μg/ml streptomycin, 2 mM L-glutamine. Jurkat E6-1 cells (ATCC) were cultured in RPMI 1640 medium supplemented with 10% (v/v) foetal calf serum (FCS), 100 units/ml penicillin, 100 μg/ml streptomycin, 2 mM L-glutamine.

### DNA constructs

The constructs LCA.1 (CD45RO) and LCA.6 (CD45RABC) were used for transfections and as templates for PCR (McKenney et al., 1995). The primers used in this study are shown in Table 3. Fragments encoding the CD45 extracellular domain (ECD) and transmembrane region were amplified by PCR and cloned into pEGFP-N1 (Clontech) using the EcoRI and BamHI restriction sites. The primers for amplification were “CD45-full length” and “CD45-antisense” using LCA.1 for pEGFP-CD45RO ECD or LCA.6 for pEGFP-CD45RABC ECD as a template. For the pEGFP-CD45 d4 construct, the primers “d4” and “CD45-antisense” and LCA.1 as a template were used. The constructs below were synthesized by PCR using pEGFP-CD45RO ECD as a template and deleting DNA encoding distinct domains of the CD45 ECD. All primers were 5’-phosphorylated. The primers used were “d2-d4-DpnI” and “antisense-DpnI” for pEGFP-CD45 d2-d4, “d3d4-DpnI” and “antisense-DpnI” for pEGFP-CD45 d3d4, “d1-sense” and “d1-antisense” for pEGFP-CD45 d1, “d1-sense” and “d1-d2-antisense” for pEGFP-CD45 d1d2, “d1-sense” and “d1-d3-antisense” for pEGFP-CD45 d1-d3, “d4-sense” and “d1-d2-antisense” for pEGFP-CD45 d1d2d4.

The expression vector pSG5-49K encoding wild type E3/49K of HAdV-D64/Ad19a has been described previously (Windheim & Burgert, 2002). The constructs below were synthesized by PCR using pSG5-49K as a template and deleting DNA encoding distinct domains of the 49K ECD. All primers were 5’-phosphorylated. The construct pSG5-R2R3 was synthesized using the primers “R2R3-antisense” and “R2R3-sense”, pSG5-R3 using the primers “R3-sense-new” and “R2R3-antisense”, pSG5-R1R3+D using the primers “R3-sense” and “R1-antisense”, pSG5-R1R3-D using the primers “R3-sense-new” and “R1-antisense”, pSG5-R1R2 using the primers “R1R2-antisense” and “R1R2-sense”, pSG5-R1 using the primers “R1-antisense” and “R1R2-sense”, pSG5-R2 using pSG5-R1R2 as a template and the primers “R2R3-antisense” and “R2R3-sense”.

For mammalian expression of the N-terminal ectodomain of HAdV-D64 E3/49K with a C-terminal HisTag, the corresponding DNA encoding residues 1-353 of 49K was amplified by PCR with primers “s49K-sense” and “s49K-antisense” and was then cloned into the pSG5 expression vector using the EcoRI restriction site. The pSG5-sec49K-His construct was then used as a template to introduce the deletion of the disordered region (residues 203-260) by PCR with the primers “R2-antisense” and “R3-sense-new” (all with 5’-P) to synthesize pSG5-sec49K-His-D.

### Expression and purification of sec49K-His using baculovirus in insect cells

For the expression of the N-terminal ectodomain of HAdV-D64 E3/49K (residues 20-353) in insects cells, the corresponding DNA was amplified by PCR with the primer “49K- BamHI” with an N-terminal signal sequence from melittin (honeybee) and with the primer “49K-EcoRI” with a C-terminal HisTag preceded by a PreScission Protease cleavage site and cloned into pFastBac^TM^Dual using the BamHI and EcoRI restriction sites. The sec49K-His-pFastBac^TM^Dual was transformed into DH10EmBacY cells and white colonies were selected for analysis. Bacmid DNA minipreps were prepared and analyzed by PCR to verify the insertion. For baculovirus production, 1×10^6^ Sf9 insect cells were transfected with bacmid DNA using Fugene (Promega) and cultivated for three days. The supernatant was collected and designated P0. 25 ml 1×10^6^ Sf9 cells/ml were infected with 3 ml of P0 and cultivated for four days. The supernatant was collected and was designated P1. An aliquot of the supernatant was analyzed for sec49K-His expression. 300 ml 1×10^6^ Sf9 cells/ml were infected with 4 ml of P1 and cultivated for three days. The supernatant was collected and designated P2. 1000 ml 3×10^6^ Sf9 cells/ml were infected with 20 ml (1:50) of P2 and cultivated for 5 days for protein expression. The supernatant was collected and the His-tagged sec49K was purified with HisTrap excel Ni beads (GE Healthcare), eluted with 300 mM imidazole and dialysed against buffer containing 50% glycerol (50 mM Tris pH 7.5, 150 mM NaCl, 0.1 mM EGTA, 50% glycerol, 0.03% Brij-35, 1 mM benzamidine, 0.1 mM PMSF). Aliquots were stored at −20°C.

### Transfection, cell lysis and immunoblotting

293T cells were transfected with DNA plasmids mixed with polyethyleneimine (Durocher et al., 2002) and extracted with lysis buffer after 48 h (50 mM Tris-HCl pH 7.5, 1 mM EGTA, 1 mM EDTA, 1% (w/w) Triton X-100, 1 mM Na_3_VO_4_, 50 mM NaF, 5 mM sodium pyrophosphate, 10 mM sodium β-glycerolphosphate, 0.27 M sucrose, 50 mM iodoacetamide, complete proteinase inhibitor cocktail (Roche)). After centrifugation for 15 min at 18,000 x g, the supernatant (termed cell extract) was decanted. Aliquots of the cell extracts (20 μg protein) were analyzed by SDS-PAGE and immunoblotting. Blots were developed with SuperSignal West Pico Chemiluminescent Substrate (Thermo Scientific) and images were generated with the LAS-3000 Luminescent Image Analyzer (Fujifilm).

### Generation of supernatants containing sec49K full-length or deletion mutants

293T cells were transfected with constructs encoding full-length 49K or deletion mutants. 24 h after transfection the medium was changed to serum-free medium and HEPES pH 7.4 was added to a final concentration of 25 mM. After six days the medium (supernatant) was collected.

### Stimulation of Jurkat T cells

5×10^6^ Jurkat T cells were serum-starved for 12-16 h and then preincubated with 1 μg/ml sec49K-His for 30 min at 37°C in 450 μl serum-free medium. Alternatively, 2.5×10^6^ Jurkat cells were preincubated with 450 μl supernatant from transfected cells containing sec49K or deletion mutants. Subsequently, the cells were stimulated with anti-CD3ε antibody (OKT3) at 3 µg/ml for different times in a total volume of 0.5 ml. Stimulation was stopped by addition of 1 ml ice-cold Dulbecco’s PBS (DPBS). Cells were spun down (250xg, 5 min), washed once with 1 ml ice-cold DPBS, spun down again (250xg, 5 min) and extracted with lysis buffer.

### Calcium flux

Calcium flux was measured with the Fluo-4 Direct^TM^ calcium Assay Kit (F10471, Thermo Scientific). The 2x Fluo-4 Direct^TM^ calcium reagent loading solution containing 5 mM probenecid was prepared according to the manufacturer’s instructions. Jurkat T cells were serum-starved for 12-16 h and 1×10^6^ cells were centrifuged and resuspended in 200 μl serum-free medium. 200 μl 2x Fluo-4 Direct^TM^ calcium reagent loading solution were added and the cells were incubated at 37°C. After 30 min sec49K-His was added at a final concentration of 1 μg/ml. After 1 h anti-CD3ε antibody (OKT3) was added at a final concentration of 3 μg/ml for 0-30 min. At different time points the cells were removed from the incubator and the fluorescence was determined by flow cytometry with a BD Accuri C6. Subsequently, ice-cold DPBS was added, the cells were centrifuged for 5 min at 250xg, lysed and analyzed by SDS-PAGE and immunoblotting.

### Affinity purification with sec49K-His and Ni-NTA agarose

Cell extract (0.5 mg) from 293T cells transfected with CD45 ECD-GFP constructs was incubated for 45 min with 1 μg sec49K-His at 4°C with shaking. 50 μl of a 40% slurry Ni-NTA agarose (20 μl beads, Macherey-Nagel) were added followed by an incubation for 45 min at 4°C with shaking. The Ni-NTA agarose beads were centrifuged for 3 min at 500xg and the pellet was washed thrice with 1 ml lysis buffer and once with 1 ml 10 mM Tris pH 7.5. The supernatant was completely removed and the samples were resuspended in SDS sample buffer and analyzed by SDS-PAGE and immunoblotting.

### Affinity purification with His-tagged GFP nanobody and Ni-NTA agarose

Cell extract (0.5 mg) from 293T cells transfected with pSG5-49K full-length or deletion constructs was incubated for 45 min with cell extract (0.5 mg) from 293T cells transfected with CD45 ECD-GFP constructs and 2 μg His-tagged GFP nanobody at 4°C with shaking. 50 μl of a 40% slurry Ni-NTA agarose beads (20 μl beads, Macherey-Nagel) were added followed by an incubation at 4°C for 45 min with shaking. The Ni-NTA agarose beads were centrifuged for 3 min at 500xg and the pellet was washed thrice with 1 ml lysis buffer and once with 1 ml 10 mM Tris pH 7.5. The supernatant was completely removed and the samples were resuspended in SDS sample buffer and analyzed by SDS-PAGE and immunoblotting.

### Flow cytometry

Flow cytometry (FACS) was carried out essentially as described (Windheim et al., 2016). 5×10^5^ Jurkat T cells or detached 293T cells transfected with the CD45 ECD-GFP constructs were incubated in 100 μl FACS buffer (Dulbecco’s PBS/ 3% FCS) with purified sec49K-His at different concentrations or supernatants containing wild-type sec49K or deletion mutants for 45 min. After three washing steps with FACS buffer, cells were incubated with 100 μl anti-49K ECD antibody for 45 min. After three washing steps with FACS buffer, Jurkat cells were incubated with 100 μl Alexa Fluor^®^ 488-labeled anti-rabbit or anti-rat antibody (2 μg/ml, Thermo Scientific) and 293T cells were incubated with 100 μl Alexa Fluor^®^ 647-labeled anti-rabbit antibody (2 μg/ml, Thermo Scientific) for 45 min. After three washing steps with FACS buffer, 5000 cells were analyzed with a BD Accuri C6 cytometer.

CD69 upregulation was determined after TCR stimulation of Jurkat T cells for 90 min with anti-CD3ε antibody (OKT3) at 1 µg/ml. Subsequently, the cells were incubated with Alexa Fluor^®^ 488-labeled anti-human CD69 antibody or an isotype control (1 µg/ml, BioLegend) for 45 min. The washing was performed as described before. Finally, 5000 cells were analyzed with a BD Accuri C6 cytometer.

### Crosslinking

293T cells were grown on 10-cm-diameter dishes and transfected with the CD45RO ECD-GFP construct. After two days the cells were detached, spun down (5 min, 150xg) and washed with 10 ml ice-cold PBS pH 8. Then the cells were spun down again and resuspended in ice-cold PBS pH 8 at a concentration of 5×10^7^ cells/ml. 100 µl (5×10^6^ cells) were then mixed with 100 µl supernatant from cells transfected with pSG5 constructs encoding 49K or deletion mutants and incubated 1 h on ice. Subsequently, the cells were washed thrice with 1 ml ice-cold PBS pH 8 to remove amine-containing culture media and proteins from the cells. Then the cells were resuspended in 190 µl PBS pH 8 at 2.5×10^7^ cells/ml. The crosslinkers BS^2^G-d_0_ (bis[sulfosuccinimidyl] glutarate-d_0_) and BS^3^-d_0_ (bis[sulfosuccinimidyl] suberate-d_0_) were dissolved in PBS pH 8 at 10 mM. 10 µl of 10 mM stock solution were added to 190 µl cell suspension for a final concentration of 0.5 mM. The cells were incubated for 2 h on ice. Subsequently, 0.8 ml ice-cold 20 mM Tris pH 7.5/150 mM NaCl was added to stop the reaction. Cells were spun down at 300xg for 3 min. Then the cells were resuspended in 1 ml ice-cold 20 mM Tris pH 7.5/150 mM NaCl and incubated 30 min on ice for quenching. Cells were spun down at 300xg for 3 min and lysed with 100 µl lysis buffer. Cell lysates were centrifuged for 15 min at 17,000xg and the supernatant (termed cell extract) was decanted. The cell extract was analysed by SDS-PAGE with 3-8% Tris-acetate-polyacrylamide gradient gels (Cubillos-Rojas et al., 2012) and immunoblotting.

### Purification of His-tagged-GFP nanobodies

The His-tagged nanobody recognizing GFP was purified as described before (Rothbauer et al., 2008; Rothbauer et al., 2006). The purified nanobody was dialysed against buffer containing 50% glycerol (50 mM Tris pH 7.5, 150 mM NaCl, 0.1 mM EGTA, 50% glycerol, 0.03% Brij-35, 1 mM benzamidine, 0.1 mM PMSF). Aliquots were stored at −20°C.

### Statistical Analysis

Statistical analysis was performed by unpaired, two-tailed t-test. Differences in means were considered statistically significant if P < 0.05. Statistical analysis was performed using Microsoft Excel or SigmaPlot 12.0 (Systat Software Inc.).

